# Analytical validation of the PROphet test for treatment decision-making guidance in metastatic non-small cell lung cancer

**DOI:** 10.1101/2023.04.20.537648

**Authors:** Ben Yellin, Coren Lahav, Itamar Sela, Galit Yahalom, Shani Raveh Shoval, Yehonatan Elon, James Fuller, Michal Harel

**Affiliations:** OncoHost LTD, Hamelacha 17 Binyamina, Israel, 3057324; OncoHost Inc., 1110 SE Cary Parkway, Suite 205, Cary, NC USA, 27518

## Abstract

The blood proteome, consisting of thousands of proteins engaged in various biological processes, acts as a valuable source of potential biomarkers for various medical applications. PROphet is a plasma proteomics-based test that serves as a decision-support tool for non-small cell lung cancer (NSCLC) patients. PROphet combines proteomic profiling using the SomaScan technology and subsequent computational algorithm. PROphet was implemented as a laboratory developed test (LDT). Under the Clinical Laboratory Improvement Amendments (CLIA) and Commission on Office Laboratory Accreditation (COLA) regulations, prior to releasing patient test results, a clinical laboratory located in the United States that employs an LDT must examine the performance characteristics concerning analytical validity. This study describes the experimental and computational analytical validity of the PROphet test, as required by CLIA/COLA. Experimental precision analysis displayed a median coefficient of variation (CV) of 3.9% and 4.7% for intra-plate and inter-plate examination, respectively, and the median accuracy rate between sites was 88%. Computational precision exhibited a high accuracy rate, with 93% of samples displaying complete concordance in results. A cross-platform comparison between SomaScan and other proteomics platforms yielded a median Spearman correlation coefficient of 0.51, affirming the consistency and reliability of the SomaScan platform. Our study presents a robust framework for evaluating the analytical validity of a platform that combines an experimental assay with subsequent computational algorithms. When applied to the PROphet test, strong analytical performance of the test was demonstrated.

## Introduction

In recent years, precision medicine has undergone a remarkable evolution by integrating experimental assays together with advanced machine learning algorithms. This synergy harnesses the power of complex data, such as genomics, proteomics and clinical data, to reveal hidden insights and construct comprehensive and robust predictive models[1]. By combining traditional experimental assays with cutting-edge machine learning algorithms, researchers can decipher intricate patterns and identify biomarkers and correlations that might otherwise remain obscured. This symbiotic approach holds the potential to revolutionize healthcare [2].

In an era marked by such technological and computational revolutions, ensuring stability and consistency of the measurements, as well as the predictive tools, is paramount. Analytical validity serves as one of the bases upon which personalized medicine stands, offering a comprehensive evaluation of the ability of a given tool to precisely and accurately detect and quantify biomarkers of interest, as well as to obtain the same model output in repeated measurements [3]. Indeed, different studies have displayed the analytical validity of predictive tools that were developed for genomic-based tests [4, 5], proteomics-based tests [6, 7], wearable devices [8] and more. Overall, the approach for such exploration involves analytical validity examination at two levels-the experimental part and the computational part [9].

Recently we have developed PROphet, a decision-making tool for first-line treatment of advanced stage non-small cell lung cancer (NSCLC) patients. The PROphet test integrates both experimental assay and computational analysis; the experimental part involves proteomic measurements obtained using the SomaScan assay, an aptamer-based proteomics platform developed by SomaLogic Inc.; the computational part is a model generated on a cohort of NSCLC patients[10]. Here we examine the analytical validity of PROphet, focusing on the two parts that comprise the test-experimental and computational. A description of the model development and the clinical validity of the test are described in Christopoulos et al[10].

The SomaScan assay is based on Slow Off-rate Modified Aptamers (SOMAmers) – chemically-modified, single stranded oligonucleotides that bind to native proteins with high affinity and specificity – and is capable of measuring approximately 7000 human proteins over a broad range of concentrations[11, 12]. The SomaScan assay has been used for biomarker discovery in multiple fields including cancer[10, 13], infectious disease[14], cardiovascular disease[7], neurological disorders[15]. The stability of the SomaScan technology was previously explored[16-18].

Here, we investigate the analytical validity of the PROphet test in OncoHost’s CLIA-certified laboratory where the testing is performed. We begin with an examination of the experimental precision of the SomaScan technology, followed by determination of the accuracy of the measurements obtained in two different sites. Then, we specifically examine the experimental precision and accuracy of the proteins used in the PROphet model. Next, we study the computational precision of the results provided by the test. Last, we compare between the expression levels obtained using SomaScan technology and the levels obtained using alternative proteomic platforms.

## Materials and Methods

### Sample collection

Plasma samples were collected from non-small cell lung cancer (NSCLC) patients either at baseline (before treatment commencement; termed here as T_0_) or before the second treatment (termed here as T_1_). Analyses were done on T_0_ samples unless mentioned otherwise. Samples were obtained as part of a clinical study (PROPHETIC; NCT04056247). Basic clinical data is found in Supplementary Table 1, including the analyses that each patient participated in. Each patient was assigned a 7-digit code. All clinical sites received IRB approval for the study protocol and all patients provided appropriate written informed consent. Blood samples were collected from each patient into EDTA-anticoagulant tubes, and the plasma was isolated from whole blood by centrifugation at 1200 x g at room temperature for 10-20 minutes within 4 hours of venipuncture (as a multi-center clinical trial, the protocol is flexible and involves a range of up to 4 hours between sample collection and plasma isolation. The effect of plasma separation on proteomic measurement was explored, showing minor effect for isolation up to 4 hr[19]). Plasma supernatant was collected and stored frozen at -80°C. Samples were shipped frozen to the analysis laboratory. The collection protocol is consistent with the SomaLogic plasma collection guidelines.

### Proteomic profiling

Plasma samples from 88 patients were profiled using the SomaScan^®^ V4.1 platform (Boulder, CO)[11, 18]. The assay simultaneously measures a total of 7596 protein targets, out of which 7288 targets are human proteins. SomaScan technology uses fluorescence to detect protein levels and the assay results are provided in relative fluorescence units (RFU). Runs were performed at two sites (OncoHost’s laboratory in North Carolina; SomaLogic laboratory in Colorado).

Plasma samples from 76 patients were also profiled using the Olink proximity extension assay (PEA; Olink Proteomics, Uppsala, Sweden) on 13 human Olink Target 96 panels (Cardiometabolic, Cardiovascular II, Cardiovascular III, Cell Regulation, Development, Immune Response, Inflammation, Metabolism, Neuro Exploratory, Neurology, Oncology II, Oncology III, Organ damage) as previously described [20]. Two samples per patient were profiled (i.e., T0 and T1 samples; see above). The assay results are provided in relative units called Normalized Protein eXpression (NPX) calculated from Ct values (as reported using real-time qPCR) following data normalization.

### Experimental precision

Intra-plate experimental precision was examined based on four technical replicates of four different patient samples, all assayed in the same running batch. Sample preparation procedure was carried on two different days by two different technicians. The intra-plate experimental precision was calculated for each technician and was quantified using Coefficient of Variation (CV). The CV was calculated for each protein and was assessed for each technician by the ratio σ⁄μ×100, where σ denotes the standard deviation of all four replicates, and μ is the mean protein level across all four replicates.

Inter-plate experimental precision was determined on four patient samples assayed in four separate runs on four different days by two technicians. Sample preparation was performed by two different technicians, and the precision was calculated per technician. For each sample, all the RFU values were measured, and CV was calculated for each protein.

### Computational precision

Computational precision is defined as the extent to which the algorithm output for a given sample is consistent between multiple runs. Computational precision was determined by the percentage of replicates that were assigned the same result by the algorithm (e.g., for a PROphet-NEGATIVE sample the consistency is given by the percentage of replicates that were labeled by the algorithm as negative). Computational precision was calculated based on four analytical runs. Sample preparation was performed by two technicians on four days. Either six or ten replicates of 14 different samples were assayed in four different runs. The computational precision was calculated for all replicates of each sample.

### Experimental accuracy

Accuracy was defined as the extent to which the protein levels measured at OncoHost’s laboratory coincide with the measurements performed by SomaLogic for the same samples. The RFU measurements acquired by SomaLogic were taken as the reference value. Accuracy rate was quantified by the percentage of accurate protein measurements. A protein measurement is accurate if the ratio RFU_OH lab_ / RFU_SomaLogic lab_ is between 0.8 and 1.2 (which is the acceptance criterion as defined by SomaLogic), where RFU_OH lab_ is the protein level measured at OncoHost’s laboratory and RFU_SomaLogic lab_ is the reference value measured by SomaLogic for the same sample. Fourteen samples were examined.

### Cross-platform comparisons of protein measurements

Comparison between the measurements obtained in the SomaScan platform and other protein measurement platforms was performed using Spearman rank correlation. The analysis was performed based on two sources: (i) 152 samples (samples from 76 patients collected at two time points-T0 and T1) for which protein expression level measurements were performed using both SomaScan and Olink PEA platforms. (ii) To expand to additional platforms measuring protein expression levels, as well as to broaden the coverage of the cross-platform analysis, external datasets with correlation comparisons were added to the analysis. Overall, 10 external datasets were added for the correlation comparison (Supplementary Table 2), covering comparison between SomaScan versus immunoassays, PEA and mass spectrometry (MS). Spearman correlation coefficients were obtained; since the correlations reported in Dammer et al.[21], were Pearson correlations, the Spearman correlation coefficient was calculated to maintain the same coefficient calculation method for all studies. Raffield and colleagues reported correlations measured in different platforms; the average data that they reported were used here (Table S2-9 of reference [22]). For cases reporting more than one value for a given protein (e.g., in the study of Kukova et al. [23], where samples were obtained before and after operation), the median level was used.

### Data analysis

Data analysis was performed using python, R, GraphPad Prism (San Diego, California, USA, http://www.graphpad.com) and Perseus computational platform[24]. For CV calculation, non-transformed RFU levels were used. Enrichment analysis was done using 1D enrichment test[25].

### PROphet model

The PROphet model provides information that can potentially aid in patient management and treatment decisions. For a thorough description of the algorithm and its clinical validity, please refer to Christopoulos et al.[10]. The model output is clinical benefit probability prediction for each patient. Based on the clinical benefit probability, the patients are assigned a PROphet-POSITIVE or -NEGATIVE result using the median clinical benefit probability as a threshold.

## Results

The PROphet test is based on proteomic measurements followed by computational analysis. As such, the analytical validation of the test reported here was investigated on two levels: (i) experimental precision and accuracy; (ii) computational precision.

### Experimental precision

Intra-plate experimental precision, which describes the variability of sample measurements in the same running batch, was determined by calculating the coefficient of variation (CV) of a set of technical replicates from a single SomaScan assay run. The median intra-plate CV ranged between 3.284% and 5.812% (overall median of 3.916%), with a median Q1 and Q3 of 2.601% and 6.053%, respectively (Fig. 1A and Supplementary Table 3A). As the sample preparation was performed by two technicians, the variability between the technicians was examined as well and displayed similar intra-plate CV measurements for the two technicians (paired t-test p-value = 0.8170; Fig. 1B).

**Fig. 1:**
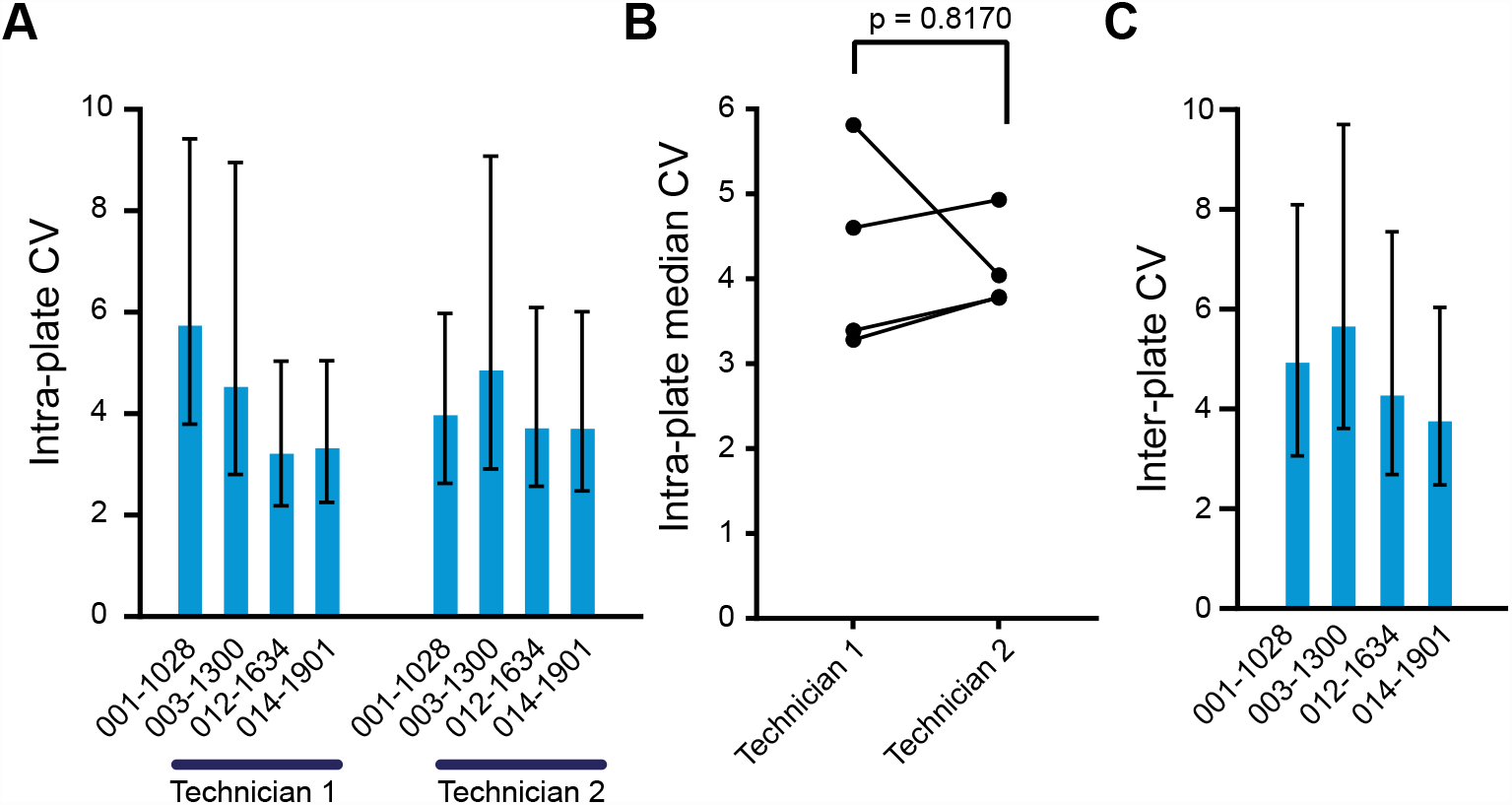
Experimental precision of all measured proteins. A. Median intra-plate CV calculated based on measurements of four technical replicates of four different samples by two technicians. The bar indicates the median and the error bars indicate the interquartile range for each sample. B. Intra-plate range comparison between technicians for all proteins. Paired t-test p-value is presented. C. Median inter-plate CV calculated based on measurements assayed in four separate runs on four different days by two technicians. The bar indicates the median and the error bars indicate the interquartile range for each sample.

The inter-plate experimental precision describes the assay’s variability between runs, days and technicians. The median inter-plate CV ranged between 3.826% and 5.731% (overall median of 4.673%), with a median Q1 and Q3 of 2.876% and 7.825%, respectively (Fig. 1C; Supplementary Table 3B), displaying similar values as the intra-plate CV. Altogether, these results demonstrate that the SomaScan assay serves as a precise proteomics platform in terms of technical replicates either within or between running batches.

### Experimental accuracy

Another important feature of a clinical test is the ability to obtain reproducible measurements between different testing sites. To determine the experimental accuracy rate, 14 patient samples were assayed in OncoHost’s laboratory by two technicians, and the results were compared to those obtained by the SomaLogic laboratory (see methods). The accuracy rate was high, spanning between 72% and 96% (Fig. 2A-B), with a median accuracy rate of 88%, without a difference between the two technicians in any of the examined samples (Supplementary Fig. 1). Overall, the results demonstrate high accuracy when comparing between two sites, with no observable difference between technicians.

**Fig. 2:**
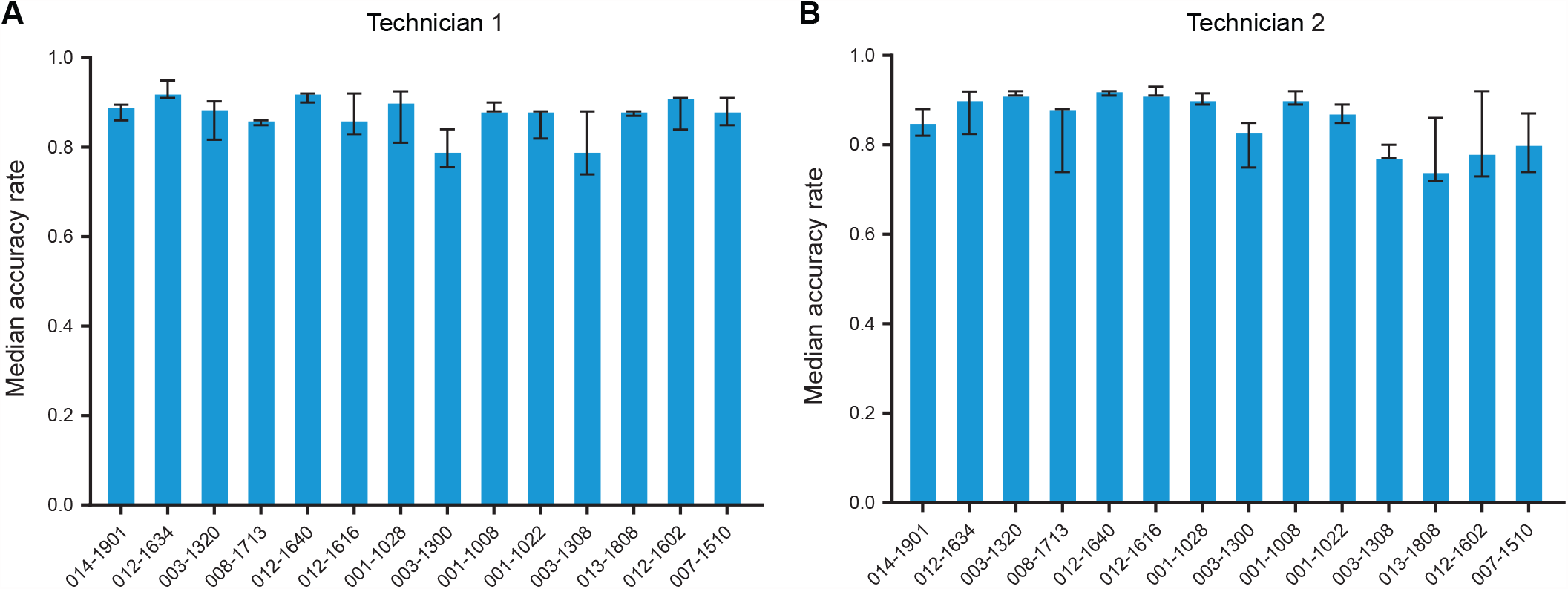
Accuracy rate of measurements between sites for all proteins. The measurements were performed at OncoHost’s laboratory and a reference laboratory (SomaLogic’s laboratory). Shown are the median accuracy rates per technician (A. and B.). The accuracy was determined based on the ratio between the measurements obtained in OncoHost laboratory relative to the reference laboratory. An accurate measurement is defined when the ratio lies between 0.8 and 1.2. The plot displays the accuracy rate (the fraction of accurate protein measurements) per sample.

### Experimental precision and accuracy of the PROphet model proteins

As the aim of the study was to assess the analytical validity of the PROphet model, we wanted to specifically examine the predictive proteins of the model [10] and to compare to the entire set of proteins measured using the SomaScan assay. One of the criteria for selecting the model proteins is their stability under variations in the measurement process, and as such they are expected to have improved precision and accuracy compared to the platform proteins as a whole.

The model proteins displayed inter-plate CV ranging from 3.018% and 4.591% (overall median of 3.474%), with a median Q1 and Q3 of 2.458% and 5.204%, respectively (Fig. 3A and Supplementary Table 4A). When comparing the intra-plate CV of the model proteins to that of the entire set of proteins, there was no significant difference in the median CV, though there was a trend towards lower CV values for the model protein set (p-value = 0.0619; Supplementary Fig. 2A), while Q3 was significantly lower in the model protein set compared to all proteins (p-value = 0.0267; Supplementary Fig. 2B). The histogram of the intra-plate CV for the entire protein set spanned over a larger range than that of the model proteins, while the latter were significantly enriched with lower CV values (1D enrichment analysis, p-value = 6.183E-05; Fig. 3B). Examination of the inter-plate precision of the model proteins revealed a significant difference when compared to all measured proteins, with a median and Q3 inter-plate CV of 3.474% and 3.867%, respectively (Fig. 3C; Supplementary Table 4B; Supplementary Fig. 2C-D; paired t-test p-value = 0.0212 and 0.0250 for the median and Q3, respectively). The range of the histogram of the inter-plate CV was larger for the entire protein set, while the model proteins were significantly enriched with lower CV values (1D enrichment analysis, p-value = 1.4496E-09; Fig. 3D). Last, the model proteins displayed higher accuracy in all samples (Fig. 4A and 4D). Altogether, the results imply that superior analytical precision and accuracy were achieved with the model proteins in comparison to all measured proteins.

**Fig. 3:**
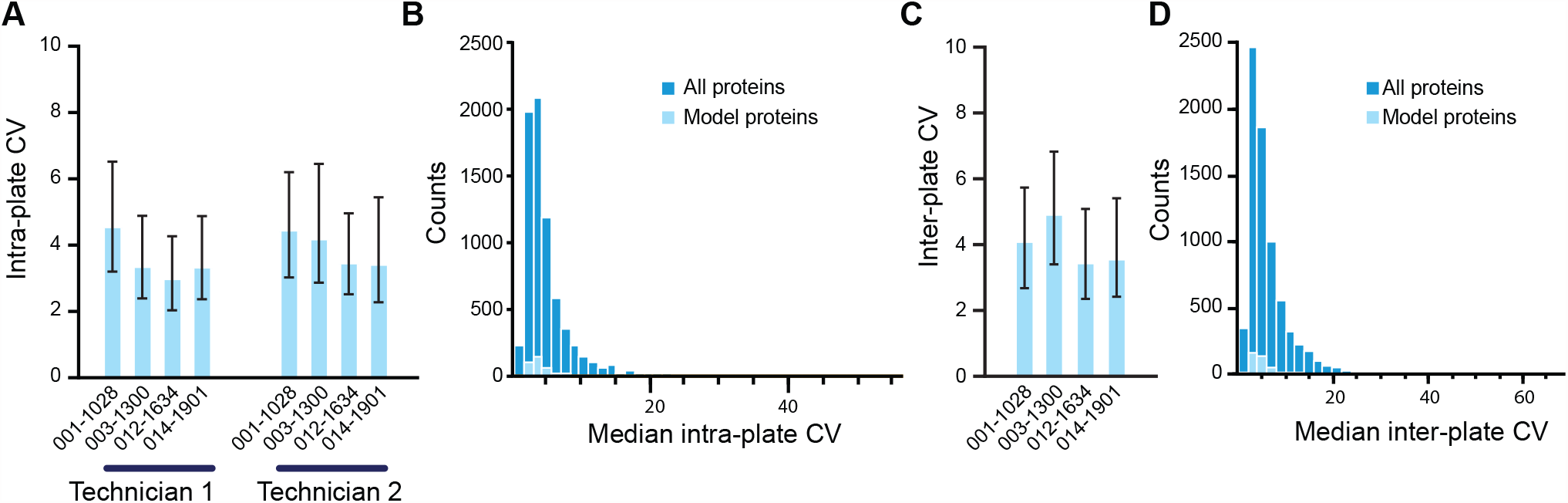
Experimental precision of the PROphet model proteins. A. Median intra-plate CV calculated based on measurements of four technical replicates of four different samples by two technicians. The bar indicates the median and the error bars indicate the interquartile range for each sample. B. Histogram of the median CV values of all proteins (dark blue) compared to the median CV values of the model proteins (light blue). C. Median inter-plate CV calculated based on measurements assayed in four separate runs on four different days by two technicians. The bar indicates the median and the error bars indicate the interquartile range for each sample. D. Histogram of the median CV values of all proteins (dark blue) compared to the median CV values of the model proteins (light blue).

**Fig. 4:**
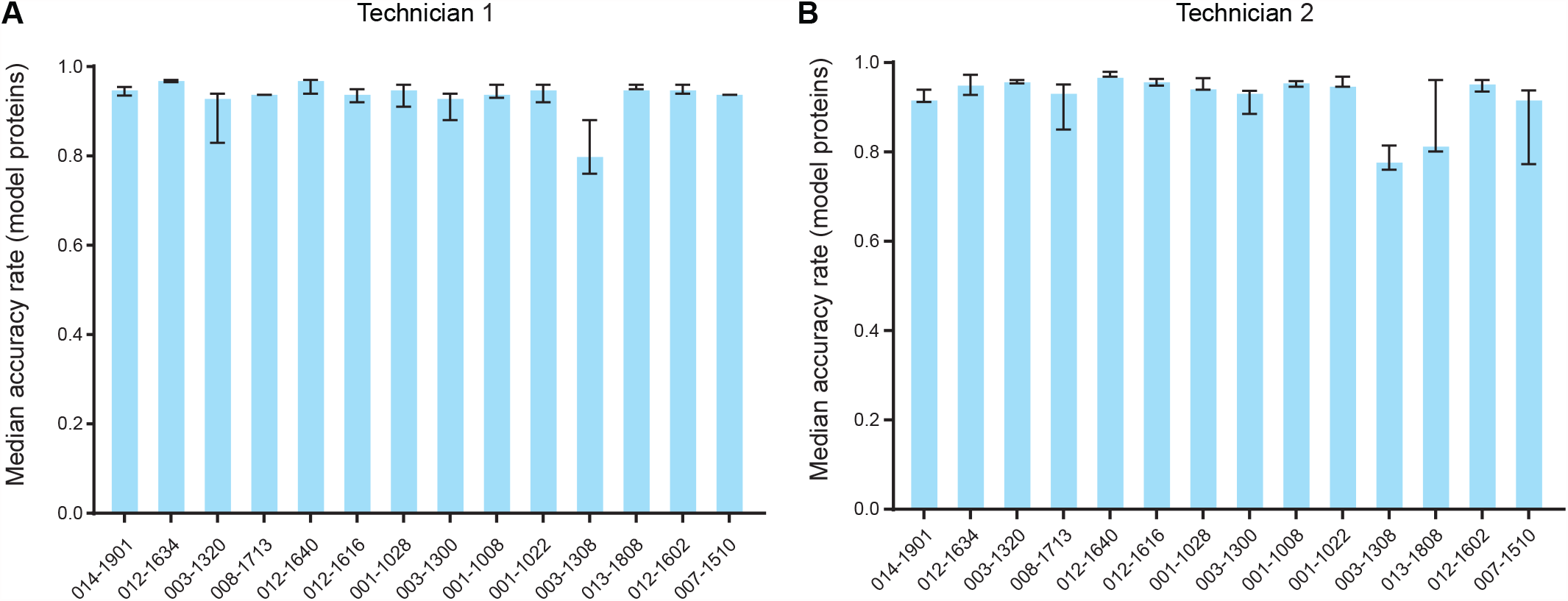
Accuracy rate of measurements between sites for the model proteins. The measurements were performed at OncoHost’s laboratory and a reference laboratory (SomaLogic’s laboratory). Shown are the median accuracy rates per technician (A. and B.). The plot displays the accuracy rate (the fraction of accurate protein measurements) per sample.

### Computational precision

The output of the PROphet model, the PROphet score, is a range between 0 and 10; Patients with a PROphet-POSITIVE result (score ≥5) survive significantly longer than patients with a PROphet-NEGATIVE result (score <5). The clinical validity of the model can be found at Christopoulos et al.[10], while here we focus on its analytical validity. Computational precision describes the consistency of a model result, PROphet-POSITIVE or -NEGATIVE in this case, when the procedure is applied repeatedly to multiple measurements of a single sample. To evaluate the test’s computational precision, we ran 14 different patient samples repeatedly in four running batches on four different days, with the addition of inter-plate experimental variability. Examination of the scaled clinical benefit probability showed that repeated measurements of 13 samples returned the same PROphet result as the reference sample (Fig. 5A-B), indicating high consistency between repeated measurements. In addition, the standard error was low for all samples, with a median of 0.1689 (Fig. 5C). Altogether, these findings imply high computational precision for the PROphet test.

**Fig. 5:**
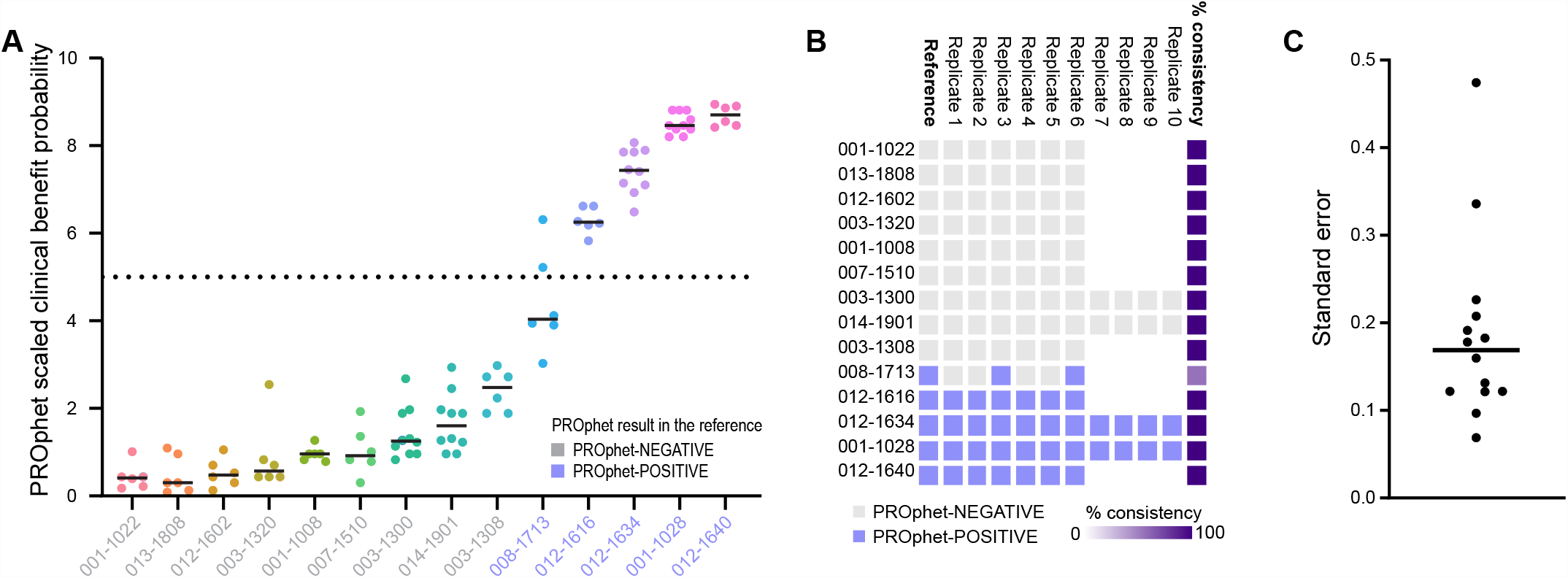
Computational precision of the PROphet model. A. The PROphet result obtained in multiple repeated measurements of the same sample. The line indicates the median of each sample. B. The consistency percentage of the algorithm to provide the same PROphet result (negative or positive). C. The standard error for the consistency.

### Cross-platform comparisons of protein measurement

Next, we sought to study the consistency between SomaScan measurements compared to other protein detection platforms. Such comparison enables an assessment and verification of protein measurements obtained using different platforms. To this end, we ran 152 samples from 76 patients in SomaScan and Olink proximity-extension assay (PEA)-based platforms. The PEA platform enabled the profiling of 1196 proteins, 1105 of which were common to the SomaScan platform. In addition, previous studies have compared between the SomaScan assay and other protein biomarker detection platforms, including traditional immunoassays, PEA and mass spectrometry (MS))[13, 21-23, 26-31]. We combined 10 such studies in order to obtain a larger coverage of proteins, as well as to add an additional source for comparison (Supplementary Table 2). Altogether, there was a Spearman rank correlation for 2247 proteins (Fig. 6). The median correlation was 0.5077, with a good agreement between correlations obtained from different studies (Supplementary Fig. 3 displays the histogram of the standard deviation for each protein, with a median standard deviation of 0.17041), though there were cases with either lower or higher correlation profiles. Notably, there were major differences in the cohort size between the different studies, which may affect the results. Immunoassays, which enable measurement of fewer proteins, varied in their correlation profile, ranging from a median of 0.1639 to 0.6875. The Spearman correlations between MS and SomaScan displayed a similar trend as was observed for the correlation profile of PEA-SomaScan, with slightly improved median correlation of 0.5366. In addition to inter-platform correlations, protein verification can be done based on cis-protein quantitative trait locus (cis-pQTL) analysis, as well as by verification of proteins using MS measurements (Supplementary Fig. 4). Examination of published data[22, 31-33] revealed 2036 proteins that were verified by at least one approach, while approximately half of them were verified by two or more studies. When examining proteins that were verified either using cis-pQTL, MS-based validation or Spearman correlation equal to or above 0.5, in total 2491 proteins remained, covering 34% of the proteome examined by SomaScan. Correlation data was available for 191 out of 388 model proteins, showing a very similar trend to the entire protein set (Fig. 6A and 6B). Comparison between the model protein set and all proteins showed no significant overall difference in the correlation coefficient distribution (Mann-Whitney test p-value for the median correlation = 0.9667). When adding cis-pQTL and MS-based verification together with model proteins with Spearman correlation above or equal to 0.5, 199 proteins remained, accounting for more than 50% of the model proteins, while 40 of which were verified by all approaches.

**Fig. 6:**
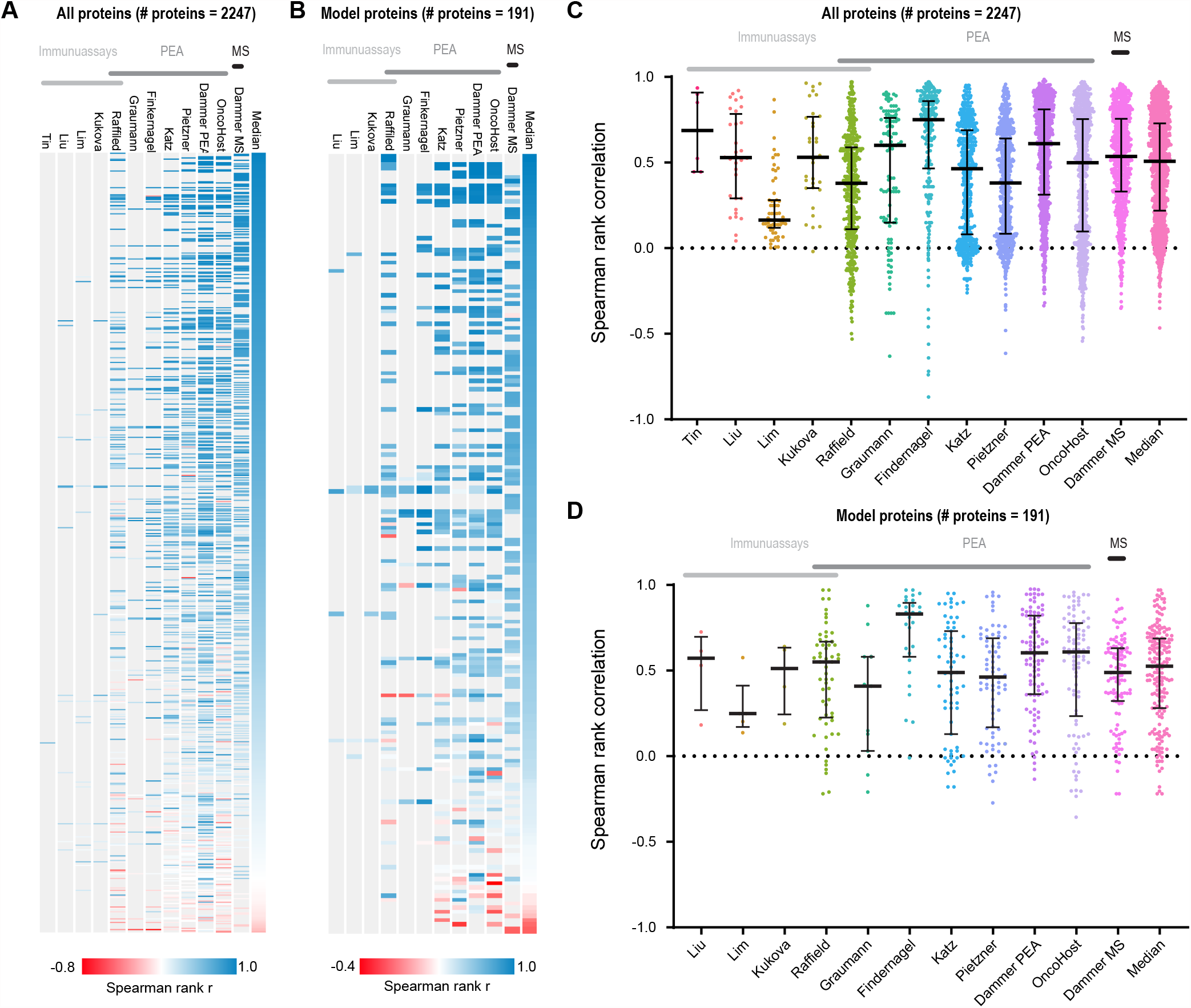
Correlations between SomaScan platform and other proteomics platforms. A. and B. A heatmap displaying the Spearman rank correlation coefficients for all proteins with available correlation data (A) and the model proteins with available correlation data (B). C. and D. The distribution of the Spearman rank correlation coefficients for all proteins with available correlation data (C) and the model proteins with available correlation data (D).

## Discussion

When developing a tool for clinical usage it is of great importance to perform a thorough assessment of the biomarker discovery platform of choice, as well as to explore its analytical validity. Here we presented the analytical validity of the PROphet tool, which is based on proteomics profiling of blood plasma followed by computational analysis.

Overall, the SomaScan platform displayed strong analytical capabilities, with low CV values within the same running batch assayed on different days, between batches and between technicians. In addition, high accuracy was observed between sites, consistent with previous reports[16-18, 27]. Altogether, these findings may be an outcome of the automatic nature of most of the sample preparation procedure. Notably, the CV values are lower than those reported for MS- and PEA-based proteomics[34, 35]. In addition, repeated measurements of the same sample displayed high consistency in terms of obtaining the same PROphet result. These properties are important not only for a clinical test, but also for the biomarker discovery process. Technologies with higher CV values require larger cohorts to overcome the inherent noise in clinical data as well as to handle the complexity of the plasma proteome; therefore, one of the advantages of SomaScan’s technology for biomarker discovery is that it performs well even with small cohorts. Notably, the PROphet model proteins displayed improved analytical properties in comparison to the entire set of proteins measured by the assay. Specifically, lower CV values and a higher accuracy rate were obtained for the model protein. This may result from the process of model protein selection during model development, where the proteomic dataset was first narrowed down to a set of proteins with high analytical reliability. In addition, the model proteins were selected iteratively using a cross-validation process, where in each iteration a different subset of patients was examined; this enabled the selection of proteins that are more confidently different between the examined groups (patients with and without clinical benefit). Clinical datasets can be complex, noisy, and contain many variables explored on a limited cohort, making it difficult to identify meaningful biomarkers that can be generalized to a new cohort of patients. Overall, the analytical superiority of the model proteins demonstrates the importance of applying good practice when identifying predictive biomarkers during model development.

Previous studies have compared between SomaScan measurements and those obtained using other proteomics platforms. Here we summarized multiple such studies and added a dataset that compares between SomaScan and PEA, together encompassing approximately 30% of the proteome examined by SomaScan V4.1. When adding protein verification using pQTL or MS analysis, along with high-correlation across platforms, our analysis covered 30% of the SomaScan proteins and 50% of the model proteins, all of which can be considered as having high-confidence agreement between platforms. Importantly, low correlation between platforms does not necessarily result from specificity or cross-reactivity issues in one of the examined platforms. There may be multiple reasons for such discrepancy, including different proteoforms examined in the different platforms or affinity differences between technologies. Overall, the specificity of the SomaScan assay arises from two main factors[18]. (i) Aptamer design procedure, where aptamers are designed to specifically bind to their target proteins with high affinity and selectivity. This is done in a process that involves screening of a large library of aptamers for those that specifically bind to the target protein. During the aptamer design process, each aptamer is characterized thoroughly, leading to the selection of very high affinity aptamers. (ii) Quality control procedures including performance testing with known protein standards and performance monitoring over time.

### Conclusion

To conclude, here we explored the analytical validity of the PROphet test, which is based on the SomaScan platform both experimentally and computationally. Overall, the test displayed strong analytical capabilities, enabling robust and reproducible predictions.

## Acknowledgements

We thank Dr. Kimberly McGregor, Kathryn Jenko and Prof. Adam Dicker for feedback and support.

## Author contribution

**Ben Yellin:** Data curation, Formal analysis, Investigation, Writing - review & editing; **Coren Lahav:** Data curation, Investigation, Writing - review & editing; **Itamar Sela:** Supervision, Writing - review & editing, Conceptualization; **Galit Yahalom:** Resources; **Shani Raveh Shoval:** Resources; **Yehonatan Elon:** Supervision, Writing - review & editing, Conceptualization; **James Fuller:** Supervision, Conceptualization; **Michal Harel:** Data curation, Formal analysis, Writing - original draft, Conceptualization.

