## Supplementary figures and images for "Analytical validation of the PROphet test for treatment decision-making guidance in metastatic non-small cell lung cancer"

### Supplementary figure 1

Supplementary Figure 1

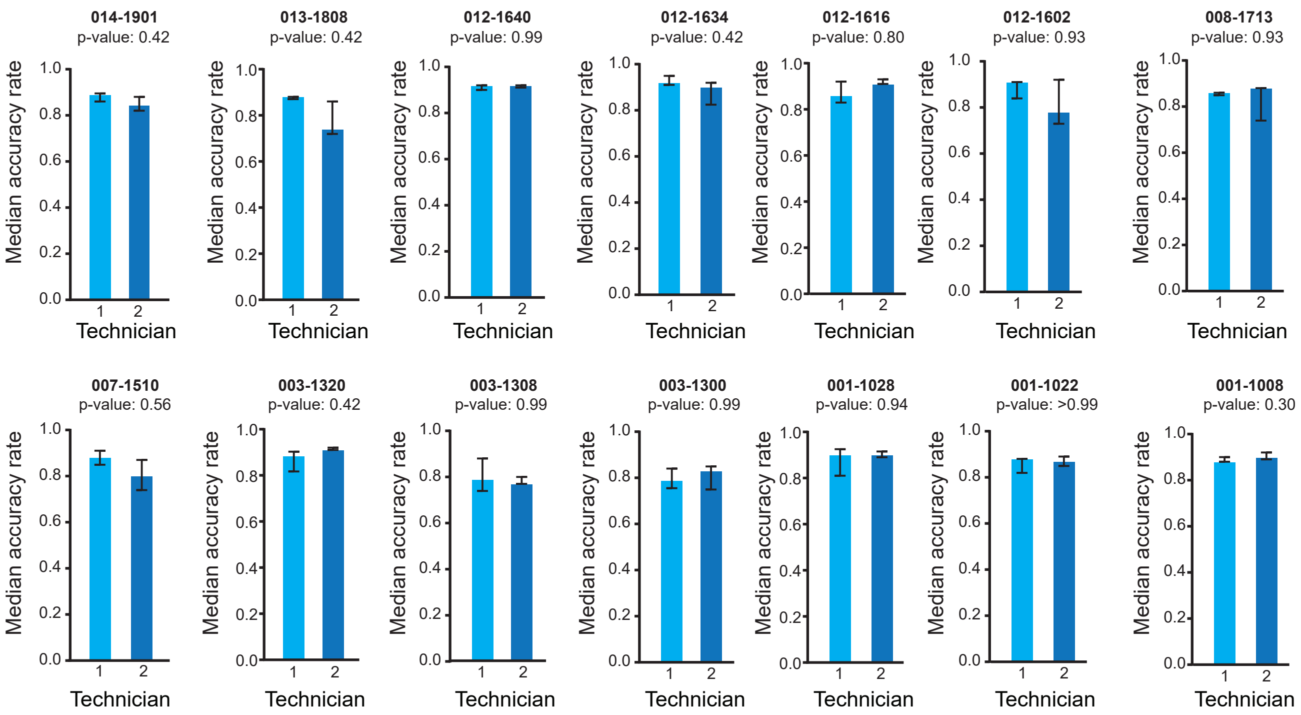

### Supplementary figure 2

Supplementary Figure 2

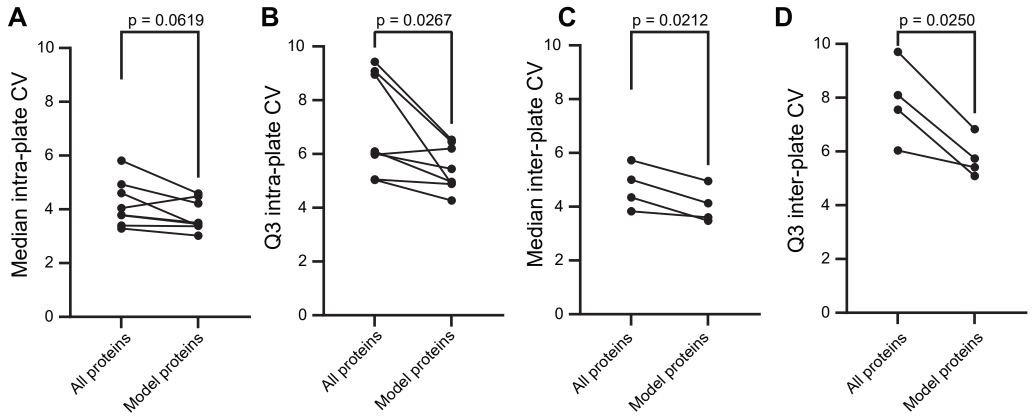

### Supplementary figure 3

Supplementary Figure 3

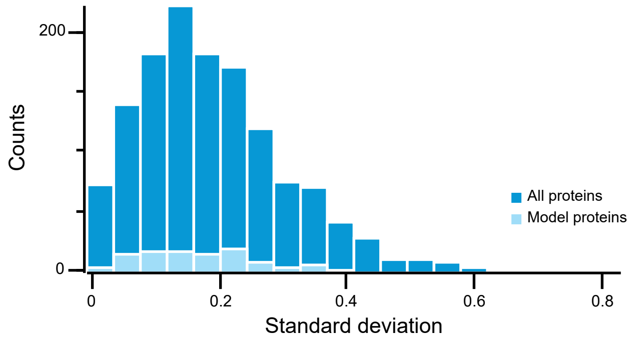

### Supplementary figure 4

Supplementary Figure 4

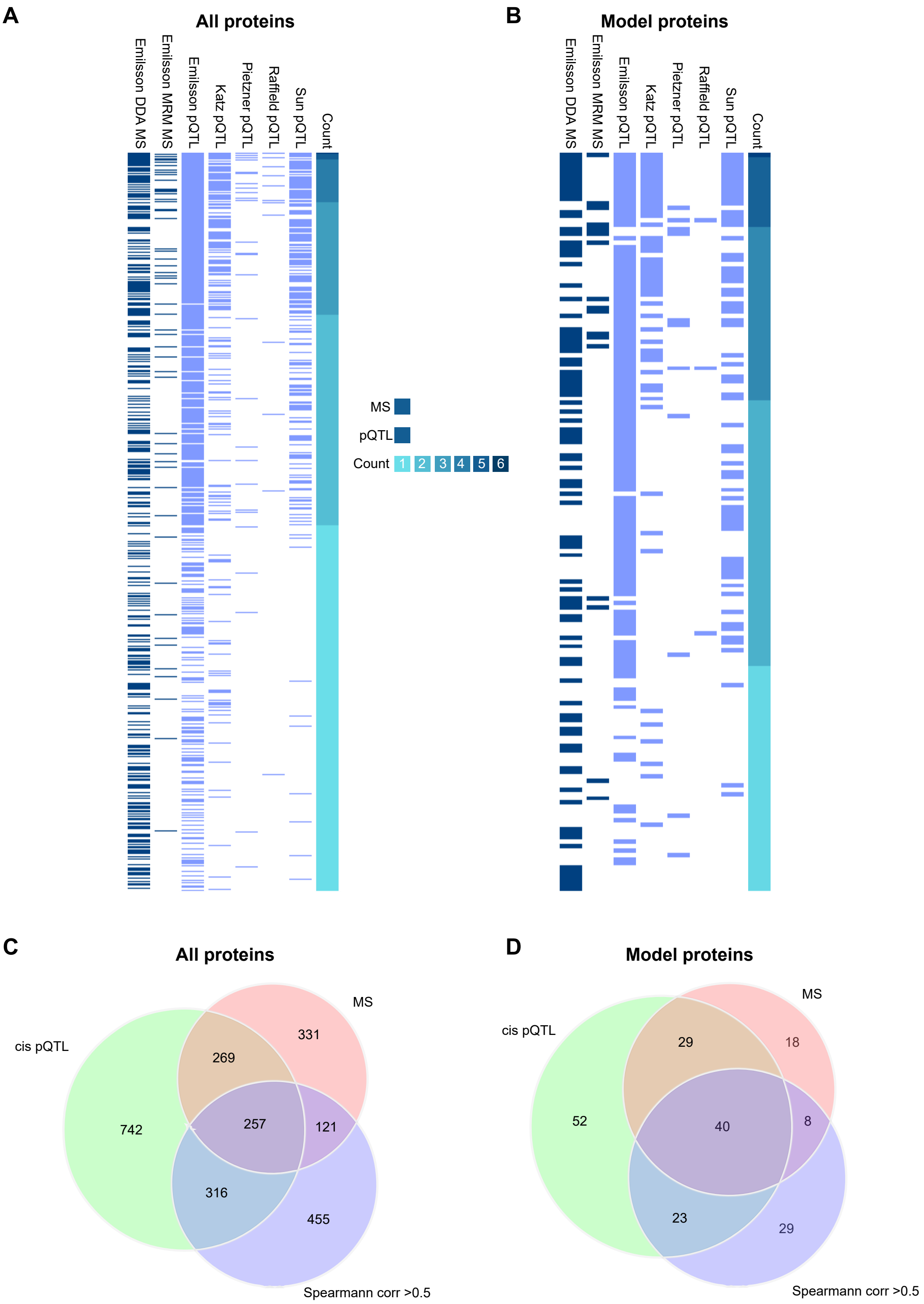
